# Mosaic loss of chromosome Y in aged human microglia

**DOI:** 10.1101/2021.11.19.469312

**Authors:** Michael C. Vermeulen, Richard Pearse, Tracy Young-Pearse, Sara Mostafavi

## Abstract

Mosaic loss of chromosome Y (LOY) is a particularly common acquired structural mutation in the leukocytes of aging men and it has been shown to correlate with several age-related diseases including Alzheimer’s disease (AD). To derive the molecular basis of LOY in brain cells, we create an integrated resource by aggregating data from 21 single-cell and single-nuclei RNA brain studies, yielding 763,410 cells to investigate the presence and cell-type specific burden of LOY. We created robust quantification metrics for assessing LOY, which were validated using a multi-modal dataset. Using this new resource and LOY-quantification approach, we found that LOY frequencies differed widely between CNS cell-types and individual donors. Among five common neural cell types, microglia were most affected by LOY (7.79%, *n*=41,949), while LOY in neurons was rare (0.48%, *n*=220,010). Differential gene expression analysis in microglia found 188 autosomal genes, 6 X-linked genes, and 11 pseudoautosomal genes, pointing to broad dysregulation in lipoprotein metabolism, inflammatory response, and antigen processing that coincides with loss of Y. To our knowledge, we provide the first evidence of LOY in the microglia, and highlight its potential roles in aging and the pathogenesis of neurodegenerative disorders such as AD.

## Introduction

A growing body of research has found that mosaic loss of chromosome Y (LOY) is an exceptionally common post-zygotic structural mutation in males^1–3^. Recent estimates utilizing the UK Biobank suggest ∼40% of men over age 70 harbor detectable LOY affecting >5% of peripheral immune cells^1^. Intriguingly, robust associations have been found between LOY and a diverse set of age-related diseases including hematologic^2, 4, 5^ and non-hematologic cancers^2, 3, 5^, macular degeneration^6^ and Alzheimer’s disease (AD)^7, 8^. Despite these correlations, the role of LOY in disease physiology and its molecular mechanisms are not well understood. Current knowledge supports the hypothesis that hematopoietic LOY events arise through mitotic missegregation errors that increase in frequency with declining systemic genomic stability^1, 9, 10^ and impaired DNA repair capabilities^1, 10, 11^. This widespread genomic dysfunction likely drives the forementioned disease associations^1, 12^. At the same time, recent high-throughput sequencing studies suggest that deletion of male-specific Y (MSY) genes and downregulation of pseudoautosomal (PAR) expression can itself lead to disease through several processes including immune system dysfunction and tumor suppressor/oncogene dysregulation^1, 13, 14^. Further investigation is required to better understand LOY and its role in disease.

Most LOY research has been conducted using readily accessible tissues like blood, buccal mucosa or tumors, and little is known about LOY occurrence and mechanisms in CNS tissue^7, 15, 16^. Nevertheless, early evidence suggests LOY does occur in the brain. Studies using WGS^16^ and qPCR^15^ both found modest but significant indications of age-dependent LOY in the dorsolateral prefrontal cortex (DLPFC). Extreme downregulation of chrY, a proxy for genomic LOY, has also been observed in multiple brain regions and was shown to increase risk of AD development^8^. Others^15^ have hypothesized that proliferative CNS cell-types such as microglia^17, 18^ and oligodendrocyte progenitor cells (OPC)^19^ are likely more prone to LOY accumulation than terminally differentiated cell-types (i.e., neurons, oligodendrocytes). Microglia are the resident macrophage and primary immune cells in the CNS that have critical roles maintaining brain homeostasis and neural function^20^. Microglial populations originate from erythro-myeloid progenitors (EMP) in the yolk-sac^21, 22^ and after colonizing the CNS are an isolated population with minimal replacement from the peripheral immune system^23–25^. Microglia also display a dramatic proliferative capacity when in an inflammatory state, plausibly leading to increasingly likely LOY-causing missegregation events during aging and neurodegeneration ^26^. However, it remains unclear if LOY detected in cortex tissue occurs locally or is derived from LOY-affected peripheral immune cells that have crossed the blood brain barrier (BBB), a process that is common during late stages of neurodegeneration^27^.

Recent studies have quantified LOY through bulk^5, 8, 13, 16^ and single-cell (sc) RNAseq^1, 13, 14^ profiling. When using the same set of samples, LOY readouts from RNAseq and array-based genotype technologies show robust pairwise correlation, providing confidence in the accuracy of each platform independently^13^. Further, recent advances in single cell multi-modal profiling have enabled deeper insight into the molecular characteristics and potential disease mechanisms of LOY cells^28–30^. For example, a CITE-seq study investigating LOY in leukocytes found simultaneous downregulation of mRNA and surface protein abundance of CD99, a gene affected by chrY deletion (located in the pseudoautosomal region (PAR) shared between chromosomes X and Y^14^). Agreement between multi-modal measurements provides further evidence that current LOY estimation methods using scRNAseq are effective and accurate.

Here, we improve on previous analytical approaches to detect and characterize cell-type specific LOY in the brain using scRNA-seq data. In total, we aggregate 21 single-cell/nuclei RNAseq datasets from 13 brain regions, including 763,410 cells from 253 individuals. Independently across five brain regions, we identify microglia as the most LOY-affected CNS cell-type. In microglial LOY populations we identify significant PAR downregulation and 213 LOY-associated differentially expressed genes. Additionally, we show agreement between multi-modal measurements of LOY using gene expression and chromatin accessibility in single cells. To our knowledge this represents the largest investigation of LOY using single-cell profiling in the brain and the first known example of LOY in microglia.

## RESULTS

### Using multi-modal single cell profiling to define LOY quantification parameters

Recent advances in single-cell assays enable simultaneous measurements of RNA and DNA, allowing for multiple lines of molecular evidence at single-cell resolution. To better understand scRNAseq LOY detection accuracy and to define informed QC thresholds for future tests, we analyzed LOY using a multi-modal ATAC and gene expression single-nuclei RNAseq dataset from B-cell lymphoma affected lymph node tissue^31^. In total, 9,380 nuclei passed QC (>1000 UMI and > 800 genes; **Methods**). Nuclei were clustered and annotated using the gene expression assay exclusively (**Fig. 1a-b**). As previously described^1, 13, 14^, the LOY status of each cell was determined using the complete lack of expression from ∼12 commonly expressed genes located in the male specific Y chromosome (MSY) region (**Methods**; **Supplementary Fig. 1**). Cells with any detectable MSY expression were classified as non-LOY/normal and those without were classified as LOY. Of all included nuclei, 23% (2250/9380) were classified as LOY (**Fig. 1c**). A vast majority of LOY nuclei (81.2%) were found in cancerous B cells (**Fig. 1d**) which was expected as LOY is common in many male neoplasms^2, 4, 5^. Next, we tested if RNA and chromatin accessibility-based estimates of LOY in the same cell would agree. To estimate LOY using scATACseq we calculated gene accessibility scores, which predict gene expression by summing chromatin accessibility counts for each gene and its promoter region (2000bp upstream). We hypothesized LOY nuclei classified using the gene expression assay would show a parallel downregulation of ATAC-based MSY gene accessibility scores. Accordingly, when using the entire dataset, we found LOY nuclei had reduced MSY normalized accessibility scores (mean = 0.010; SD = 0.0350) compared to normal nuclei (mean = 0.053; SD = 0.0527; Wilcoxon *P<*0.001; **Fig. 1f**). This was a chrY specific pattern and was not observed on other chromosomes (**Supplementary Fig. 2**). Similar trends were observed in isolated B tumor nuclei (**Fig. 1e-f**). In total, 91.4% of LOY calls determined by gene expression overlapped with those determined by ATAC, and 45.7% of all calls were shared between both technologies. Furthermore, as we selected for nuclei with more sequenced transcriptomic data (UMI counts) the agreement between the two assays increased and MSY accessibility scores in LOY nuclei approached 0 (**Fig. 1g)**. For example, when we only retained nuclei with >3000 UMI, MSY gene accessibility scores from non-LOY nuclei were 0.001 (SD = 0.011; *n*=1232), while non-LOY MSY accessibility scores remained stable (mean = 0.047; SD = 0.044; *n*=2602). For this reason, subsequent analyses estimating LOY frequency used UMI thresholds of 3000 and feature thresholds of 1000 (**Fig. 1g**). Further, consistent trends were observed for each detected MSY gene (**Fig. 1h**), which were visualized individually for ZFY and UTY (**Supplementary Fig. 3**). In each case, accessibility peaks in the LOY group were largely absent although small, noisy peaks were observed, likely representing errantly classified LOY nuclei and sequencing noise. Together, these findings support previous assumptions ^13^ that increasingly stringent UMI and feature thresholds reduce false-positive LOY calls as more transcriptional information is available and the effect of stochastic chrY gene dropout as a confounding factor is reduced.

**Fig 1.**
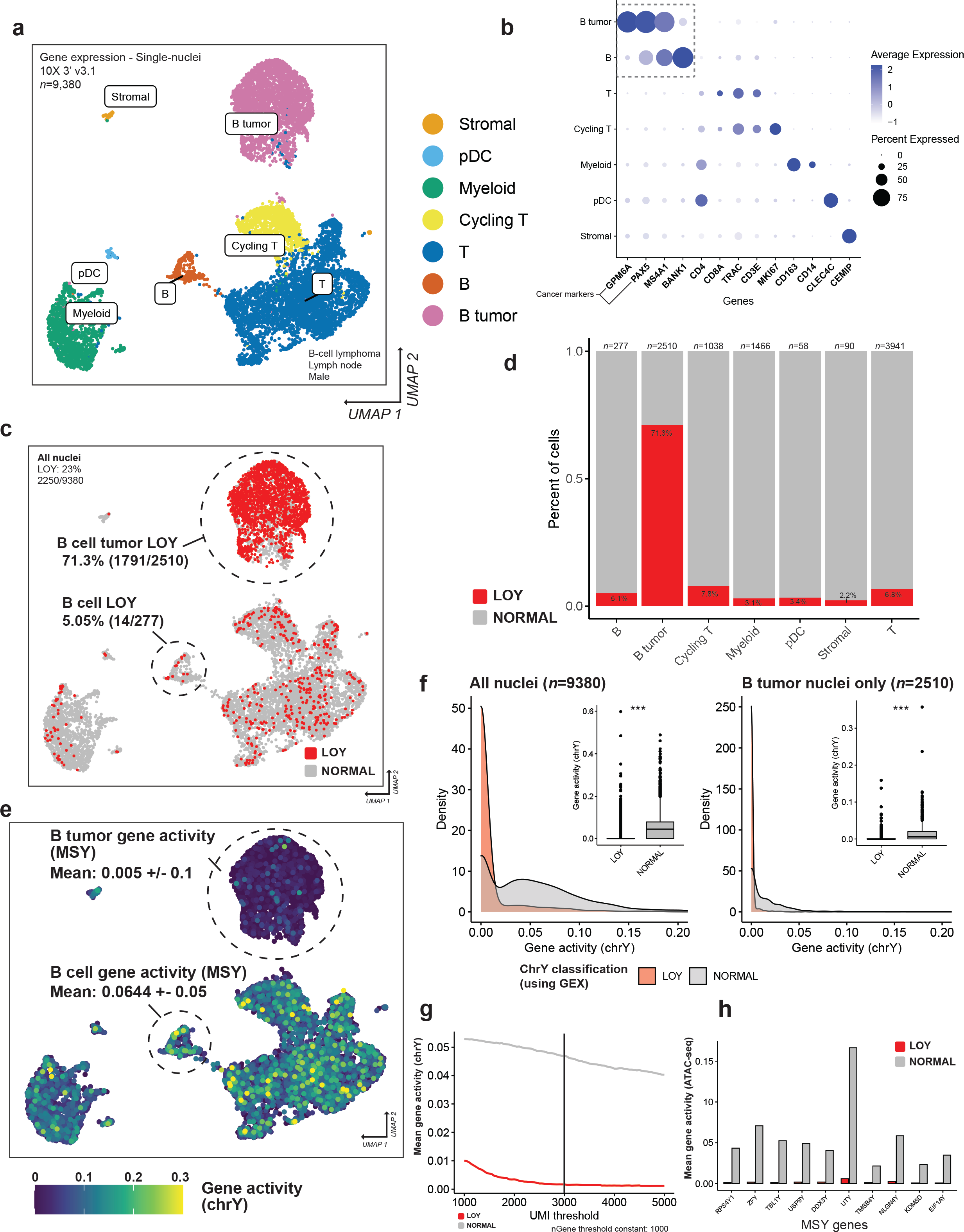
Expression-based classification of LOY agrees with ATAC-seq in the same nuclei. **a**, UMAP plot and cell-type identities of 9,380 nuclei from a male patient with non-Hodgkin’s lymphoma. Nuclei were clustered using the transcriptome. **b**, Dot plot showing canonical marker genes used to annotate Leiden-based clusters with cell-types. GPM6A and PAX5 were used to differentiate cancerous and non-cancerous B cells. **c**, UMAP plot displaying LOY nuclei (red) and non-LOY/normal nuclei (grey) which are labelled based on the expression, or lack of expression of male-specific Y (MSY) genes. **d**, Stacked bar plot showing the LOY frequencies within each cell type. LOY nuclei (red) were most frequent in cancerous B-cells (71.3%; LOY 1791/2510). **e**, UMAP plot colored by ATAC-seq derived gene-activity scores of male-specific Y genes. ATAC-seq and RNAseq were performed for each nucleus. B-cell tumor nuclei have a substantially decreased chrY gene activity in relation to other cell-types. **f**, Density plots, and boxplots (inset) comparing mean MSY ATAC gene activity of LOY (red) and normal (grey) nuclei using (left) all nuclei (n=9380) and (right) B-cell cancer nuclei only (n=2510). Significant differences between LOY and non-LOY groups were assessed using the non-parametric Wilcoxon test, resulting in the p-values shown for each comparison. All tests were one-sided. The boxes represent the 25th percentile, median, and 75th percentile. The whiskers extend to the furthest value that is no more than 1.5 times the inter-quartile range. **g**, Line plot showing mean MSY mean gene activity (ATAC-seq) of LOY and non-LOY/normal nuclei as the minimum UMI count threshold is increased. **h**, Barplots showing mean gene activity of each detected MSY gene for LOY (red) and non-LOY/normal nuclei (grey). ns P>0.05, *P < 0.05, **P < 0.01, ***P < 0.001

### Microglia show elevated LOY frequencies

To characterize cell-type specific LOY frequencies in the brain we annotated 21 publicly available single-cell (sc-RNAseq) and single-nuclei (sn-RNAseq) datasets (**Supplementary Table 1**). This yielded 763,410 male cells/nuclei from 253 donors and covered a diverse set of brain regions, cell-types, and pathologies. After filtering cells and individuals for UMI, gene expression counts, and cell-type representation (see **Methods**), 536,822 cells/nuclei were included for LOY analysis from 221 donors (242 samples), from 21 brain datasets and 13 brain regions (**Fig. 2a**). The pathologies represented included Alzheimer’s (AD; *n* = 49), Parkinson’s (PD; *n* = 9), other dementias (*n* = 16), Huntington’s (HD; *n* = 7), and ALS (*n* = 11). Age data was available for 194 donors (mean age = 59.0, age range = 0-96). See **Supplementary Fig. 1** and **Methods** for further details.

**Fig 2.**
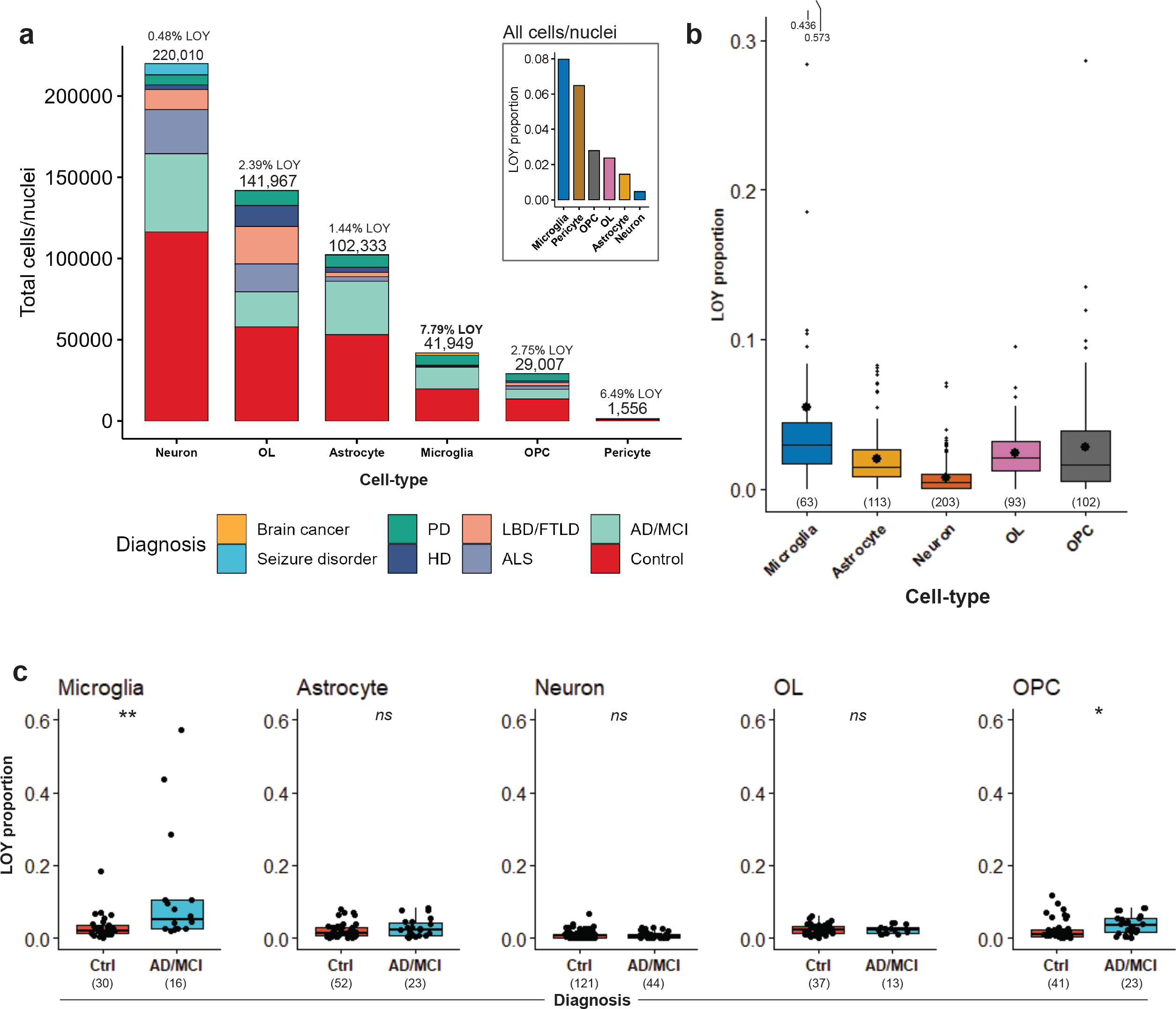
LOY frequency analysis using 505,325 cells/nuclei from 21 brain datasets. **a**, Main, stacked barplot shows total sample size of included cells/nuclei annotated to each major brain cell-type and colored by neurodegenerative diagnosis. Above each bar is the total nuclei/cell sample size and LOY frequency. Only datasets sequenced using 10X Genomics library construction were considered for the study. Cells/nuclei with greater than 3000 UMI counts and 1000 genes were included for LOY frequency analysis. Each cell-type population was considered in each subject if >75 cells/nuclei were present and MSY was commonly expressed in non-LOY cells (Methods). Inset, total LOY frequencies for each cell-type across the entire dataset. **b**, Boxplot of mean LOY frequencies of each subject across cell-types. The boxes represent the 25th percentile, median, and 75th percentile. The whiskers extend to the furthest value that is no more than 1.5 times the inter-quartile range. Large dots represent the mean. For visualization purposes, the Y-axis range was limited, and two microglia outliers were not visualized. Reference to these outliers extending off the plot (0.436 and 0.573) are included at the top and are not to scale. Subject sample sizes are provided in parentheses under each box. **c**, Comparisons of LOY frequency between AD/MCI and control samples for each cell-type. Significant difference between groups was assessed using an unadjusted, non-parametric Wilcoxon test, resulting in the p-values shown for each cell-type. All tests were two-sided. AD Alzheimer’s disease, MCI mild cognitive impairment, PD Parkinson’s disease, HD Huntington’s disease, ALS Amyotrophic lateral sclerosis, LBD Lewy body dementia, FTLD Frontotemporal lobar degeneration, OL oligodendrocyte, OPC oligodendrocyte progenitor cell, LOY mosaic loss of chrY. ns P>0.05, *P < 0.05, **P < 0.01, ***P < 0.001.

We found that LOY cells represented 1.89% (n=10,158) of the combined dataset and were observed in 199 of 242 samples (**Supplementary Table 2**). We next quantified LOY proportions across five major brain cell-types that were robustly captured. Of 41,949 tested microglia, 7.79% were classified as LOY, greater than OPC (2.75%, *n*=29,007), OL (2.39%, *n*=141,967), astrocyte (1.44%*, n*=102,333) and neurons (0.48%, *n*=220,010; **Table 1**). On average, each individual subject displayed a mean microglial LOY frequency of 5.46% (range=0-57.33%, *n*=63; **Fig. 2b**). In OPCs, OL, astrocytes, and neurons mean donor LOY frequencies were 2.75% (0-28.61%, *n*=102), 2.39% (0-9.5%, *n*=93), 1.94% (0-8.12%, *n*=113), 0.76% (0-7.1%, *n*=203), respectively (**Fig. 2b**). In microglia and OPC we also found increased LOY frequency in AD donors compared to controls (**Fig. 2c**), however, we note the tight correlation between AD and age which could explain this association. These findings suggest that proliferative CNS cell-types could also be prone to LOY accumulation.

**Table 1.** Observed LOY frequencies across CNS cell-types using scRNAseq. LOY frequencies were estimated across 21 datasets. LOY Loss of Y, OL Oligodendrocyte, OPC Oligodendrocyte progenitor cell.

| Cell type | Datasets | Donors | LOY cells | Non-LOY cells | Total cells | Total LOY proportion | Mean donor LOY | Donor LOY range | Extreme LOY (>15%) |
| --- | --- | --- | --- | --- | --- | --- | --- | --- | --- |
| Astrocyte | 16 | 113 | 1474 | 100859 | 102333 | 1.44% | 1.94% | 0-8.13% | 0 |
| Microglia | 13 | 63 | 3345 | 38604 | 41949 | 7.97% | 5.46% | 0-57.33% | 4 |
| Neuron | 19 | 203 | 1051 | 218959 | 220010 | 0.48% | 0.76% | 0-7.1% | 0 |
| OL | 10 | 93 | 3388 | 138579 | 141967 | 2.39% | 2.38% | 0-9.5% | 0 |
| OPC | 15 | 102 | 799 | 28208 | 29007 | 2.75% | 2.72% | 0-28.6% | 1 |
| Pericyte | 4 | 11 | 101 | 1455 | 1556 | 6.49% | 7.27% | 2.5-17.7% | 1 |
|  |  |  | 10158 | 526664 | 536822 | 1.89% |  |  |  |

### Transcriptional impact of LOY

We next performed differential expression (DE) tests between LOY and non-LOY cells, first as a secondary validation for LOY estimation and second to identify transcriptional events that help explain LOY processes. In particular, the diploid PAR regions (PAR1 and PAR2) are shared between the distal ends of chromosome X and Y and a whole chromosome LOY causes loss of heterozygosity (LOH) of the region leading to downregulation (**Fig. 3a**). Thus, PAR downregulation represents an independent transcriptional biomarker of LOY, as LOY is directly classified using null chrY expression. Interestingly, significant PAR downregulation was solely observed in microglia (Hypergeometric test, FDR < 0.1) which was consistent across three microglia datasets (**Fig. 3b**) covering four diverse brain regions including the somatosensory cortex, entorhinal cortex, dorsolateral prefrontal cortex, and substantia nigra (Hypergeometric test, FDR < 0.1; **Supplementary Table 3**). As a result of robust PAR downregulation in LOY populations, the remainder of the analysis was primarily focused on microglia.

**Fig 3.**
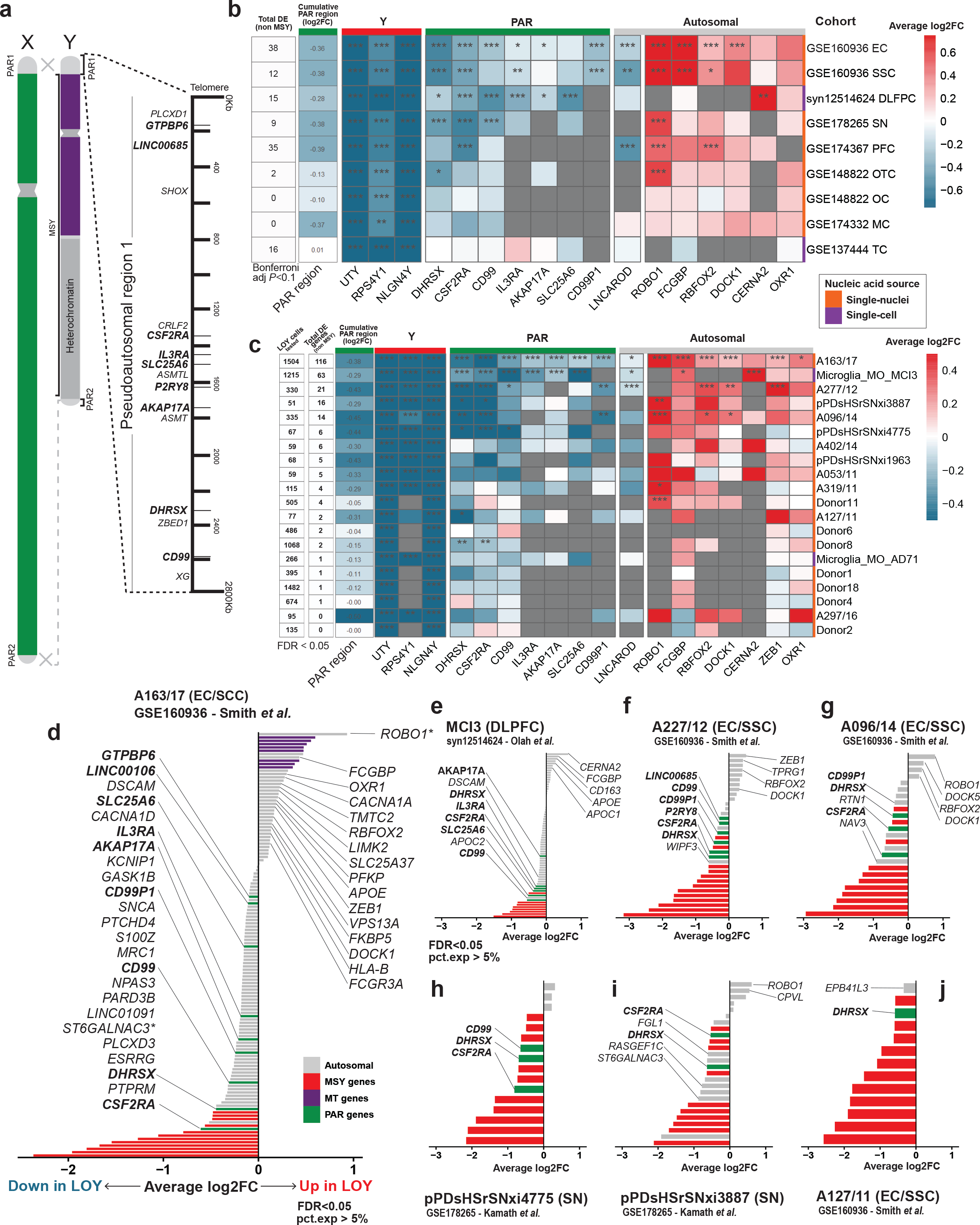
PAR region genes are recurrently downregulated in microglial populations with substantial LOY. **a**, Schematic describing the genomic context and genes residing within the pseudoautosomal regions (PAR). The PAR regions are homologous sequences shared between the X and Y chromosomes. Genes within PARs are diploid and are inherited like autosomal genes. **b**, Heatmap illustrating microglia DE (average logFC) across several single-cell/nuclei cohorts with > 100 LOY cells/nuclei. Grey values represent NAs as the gene was not expressed in >5% of cells/nuclei. If multiple brain regions were sampled within a given dataset, they were split by region and labelled accordingly. The MAST algorithm was used and sample, UMI count, detected gene count, and mitochondrial percent were regressed out. The first column on the left displays the total number of significant, non-MSY DE genes (Bonferroni P<0.1) detected per dataset. The second column displays cumulative logFC of all expressed PAR genes. Rows are coloured based on source of nuclei acid (whole cell or nuclei). Significance is provided within each cell; ns P>0.1, *P < 0.1, **P < 0.01, ***P < 0.001 (Bonferroni multiple-test correction). **c**, Heatmap illustrating microglia DE (average logFC) for individual subjects with > 50 LOY cells/nuclei. The first column on the left displays the total number of tested LOY cells/nuclei, while the second column shows the number of significant, non-MSY DE genes (FDR<0.05) detected per subject. Significance is provided within each cell; ns FDR>0.05, *FDR < 0.05, **FDR < 0.01, ***FDR < 0.001 (False-discovery rate). **d-j**, Bar plots illustrate the average log fold change (logFC) of differentially expressed (DE) genes (FDR < 0.05) between LOY and non-LOY cells/nuclei within microglia from seven subjects including: (**d**) A163/17 (AD; GSE160936), (**e**) MCI3 (MCI; syn12514624), (**f**) A277/12 (AD; GSE160936), (**g**) A096/14 (AD; GSE160936), (**h**) pPDsHSrSNxi4775 (PD; GSE178265), (**i**) pPDsHSrSNxi3887 (PD; GSE178265) and (**j**) A127/11 (Control; GSE160936). Cells/nuclei were included for DE analysis if they had > 1000 UMI counts and >800 detected genes. Select gene names are shown, and PAR gene names are labelled in bold. Autosomal, PAR, male-specific chrY, and mitochondrial gene bars are colored grey, green, red and purple, respectively. Several PAR genes are recurrently downregulated across the selected subjects, and PAR gene CSF2RA is significantly downregulated in 6/7.

Focusing our analysis on 20 donors with sizeable microglial LOY populations (>50 LOY cells) with detected PAR dysregulation, we identified, 213 unique non-MSY, microglial LOY Associated Transcriptional Events (mLATE) (mLATE) genes (FDR<0.05), including 26 autosomal mLATE genes and 7 PAR genes (CD99, CD99P1, DHRSX, IL3RA, CSF2RA, AKAP17A, SLC25A6) that were dysregulated in multiple subjects (**Supplementary Table 4; Fig. 3c**). mLATEs from seven individuals showed significant overrepresentation of the PAR (Hypergeometric test, FDR < 0.1; **Fig 3d-j**; **Supplementary Table 5-6**). Of donors with significant PAR downregulation, 6/7 were diagnosed with either AD, PD or mild cognitive impairment (MCI).

Recurrent autosomal mLATE genes displayed diverse functions in CNS development and cancer promoting processes, including roles in axon guidance (ROBO1, DOCK1, NAV3), immune cell and glioma migration (ROBO1), alternative splicing regulation and estrogen transcription (RBFOX2), phagocytosis (DOCK1, CD163), neuroinflammation (ZEB1), ganglioside biosynthesis (ST6GALNAC3) and lipoprotein metabolism (APOE). ROBO1 (Roundabout Guidance Receptor 1), part of the SLIT/ROBO signaling pathway, was the most overexpressed mLATE gene (avg logFC = 0.68; range = 0.11–1.26) and was significantly upregulated in five subjects. In our meta-analysis, SLIT/ROBO genes were rarely expressed in non-LOY microglia in any brain dataset (**Supplementary Fig. 4**), and ROBO1 was the sole SLIT/ROBO gene differentially expressed in LOY microglia. ROBO1 and the SLIT/ROBO pathways are interesting targets for future LOY studies especially because LOY has been observed in ∼20% of male glioblastomas (GBM)^33^.

Pathway enrichment analysis (Methods) identified 27 significantly overrepresented pathways (**Supplementary Table 7**) and GO terms (**Supplementary Table 8**; FDR<0.05), including lipoprotein metabolism, inflammatory response, regulation of cytoskeleton, and hemostasis (**Supplementary Fig. 5**). Although more experimental investigation is required, these results suggest that LOY microglia are likely more activated than non-LOY microglia. Inflamed, phagocytic microglia clearing injured tissue, amyloid beta and other debris proliferate rapidly, potentially leading to more frequent LOY events. We also compared our mLATEs with 489 LATEs observed in the leukocytes of aging men^13^ and found significant overlap (Fisher’s exact test *P*=0.0051). Overlapping autosomal LATE genes included B2M, FXYD5, IFITM3, ITGAX, KLHL6, RPLP0, S100Z, SCMH1, TMEM176B, and TMEM71, again highlighting roles in immune function, chemotaxis, and inflammation.

### LOY is observed across all microglia subtypes

To further investigate LOY in microglia we focused on seven datasets containing subjects with elevated LOY (>10% LOY)^34–40^ (**Supplementary Table 3**). Microglia were isolated, reclustered and annotated for microglia subtypes (**Supplementary Fig. 6-12**). Smith *et al.*^34^ (referred to as ‘Smith’) enriched for microglia and astrocyte nuclei from the entorhinal (EC) and somatosensory cortex (SSC) of AD diagnosed and control donors, while Olah *et al.*^35^ (‘Olah’) and Mancuso *et al.*^36^ (‘Mancuso’) enriched for microglia from DLPFC and temporal cortex, respectively. Gerrits *et al*.^37^ (‘Gerrits’) enriched for microglia and astrocytes from the occipital cortex. These cell-types are often underrepresented in CNS single-cell RNAseq datasets ^41^. Kamath^40^, Morabito^39^ and Pineda^38^ consisted of unsorted nuclei from the substantia nigra, DLPFC and primary motor cortex, respectively.

In the ‘Smith’ dataset, 48,748 QC’d nuclei were retained including 32,122 astrocyte (65.9%), 13,222 microglia (27.1%), and 1,304 neuronal nuclei (2.67%) from 4 AD and 4 control donors, all with relatively high per cell UMI counts (median = 6,386; IQR = 6,788.7), gene counts (median = 2,758; IQR = 1824.7), sufficient nuclei sample size (mean nuclei per sample = 3,585.1; SD = 1,439.2), and robust Y chromosome expression in biological replicates, providing a quality opportunity to compare LOY proportions in glial cells between AD and control samples. (**Fig. 4**; **Extended Fig 1**; **Supplementary Fig. 13**). Consistent with our global LOY analysis, across all ‘Smith’ samples and brain regions LOY nuclei were most common in the microglia (2,068 / 13,222; 15.64%; range (donor) = 1.65%–61.91%; **Fig. 4b**) and rare in neuronal (10 / 1,304; 0.76%; range = 0%-1.12%) and astrocyte (224 / 32,122; 0.69%; range = 0.14%-4.33%) populations. Endothelial cells (51 / 495; 10.3%), OPCs (30 / 362; 8.28%), pericytes (48 / 679; 7.06%) and oligodendrocytes (9 / 157; 5.73%) also showed elevated LOY rates as well, however sample sizes were considerably smaller and more prone to variation. Strikingly, microglial LOY frequency was substantially greater in AD donors (28.0%) compared to controls (2.76%); a trend that was consistent across both SSC and EC brain regions (**Fig. 4b**). In SSC tissue, 21.1% of AD microglia nuclei were classified as LOY compared to 1.95% in controls (**Fig. 4f-g)**, and in the EC LOY frequency was also greater in AD (32.5%) than control (3.21%) (**Fig. 4c-d**). These trends were not observed in either astrocytes or neuronal nuclei where LOY proportions were similar between both AD diagnosis and brain regions (**Extended Figure 1c-d, f-g**). Elevated LOY in the EC of AD donors is of interest as evidence suggests the EC is uniquely prone to proteopathies^42^ and is thought to be the first brain region affected by AD^43, 44^. LOY microglia were observed in every subject but were primarily seen in microglia populations of 4 samples from 2 AD donors: A096 (Age: 87; SSC LOY: 42.2%; EC LOY: 47.2%) and A163 (Age: 83; SSC LOY: 46.1%; EC LOY: 61.9%; **Supplementary Fig. 14**). These 2 subjects harbored 76.1% of the LOY population in the ‘Smith’ dataset.

**Fig 4.**
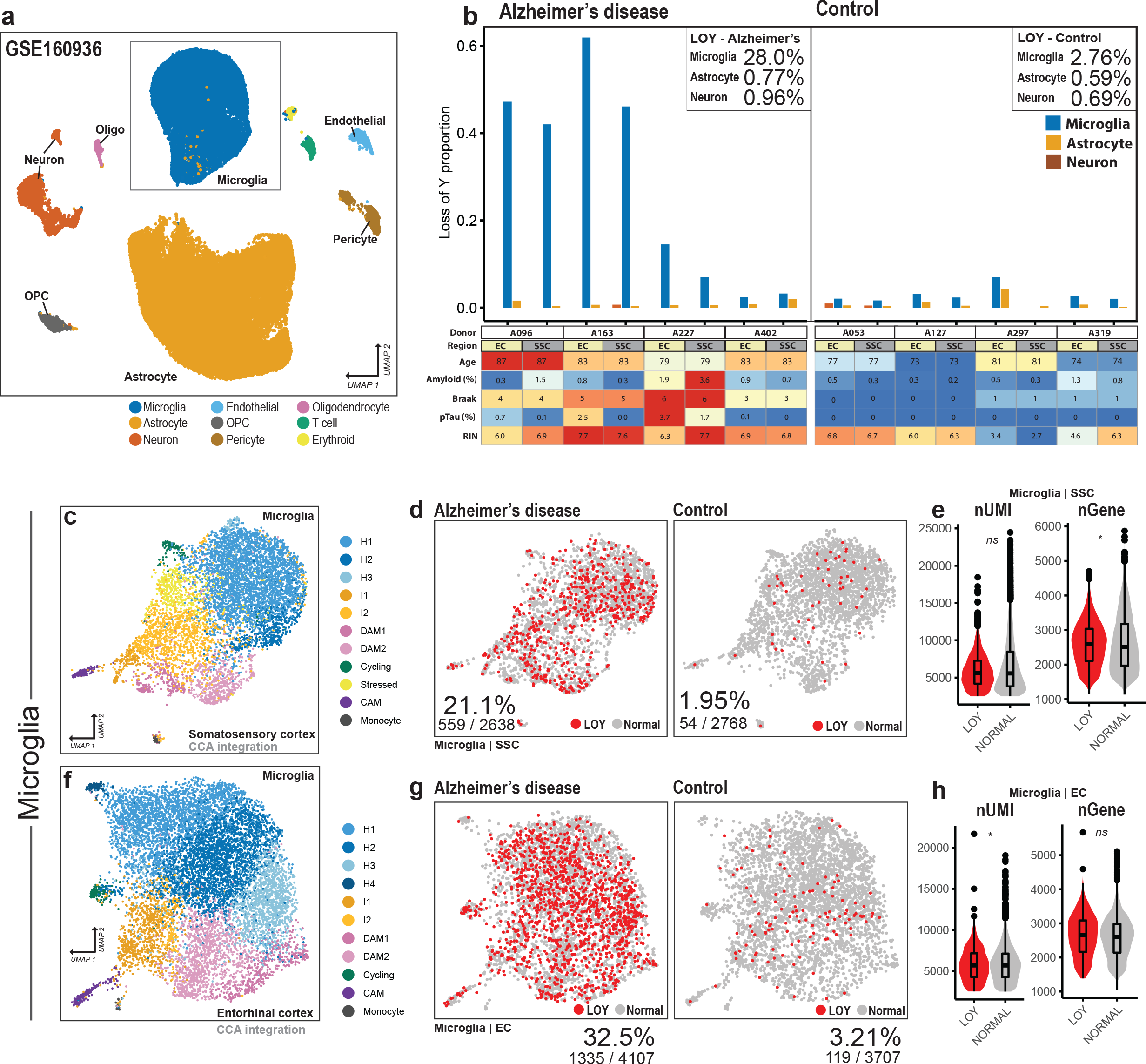
LOY frequencies are enriched in AD microglia across the EC and SSC (GSE160936). **a**, UMAP reduction and cell-type annotations of 48,748 single-nuclei brain transcriptomes from GSE160936. Markers used to annotate clusters include C3 and CSF1R (microglia), AQP4 and GFAP (astrocyte), RBFOX3 and GAD1 (neuron), and MBP (oligodendrocyte). All marker genes used to annotate clusters are available in Supplementary Fig. 9. **b**, Top, coloured bars represent LOY frequency in each sample for microglia (blue), astrocytes (orange) and neurons (red-brown). Donors are ordered by AD diagnosis, AD (left) and non-disease control (right). Each donor was sampled from both the entorhinal cortex (EC) and somatosensory cortex (SSC). Bottom, heatmap displaying additional phenotypic information of each donor/sample. Provided measures of amyloid %, Braak score and pTau % were all determined using quantitative image analysis. Microglia nuclei from SSC (c) and EC (f) were isolated, CCA integrated, and clustered. **c,f**, Subclusters were annotated using microglia gene panels (Supplementary Table 11). Homeostatic subclusters (H) are colored in various blues, while inflammatory subclusters (I) are colored in oranges and disease associated microglia (DAM) clusters are colored in magentas. **d,g**, UMAP reductions of SSC and EC split by AD diagnosis; Nuclei are colored by LOY (red) and non-LOY/normal (grey). LOY nuclei frequencies are provided at the bottom of each UMAP plot. **e,h**, Violin boxplots showing UMI counts (nUMI) and detected genes (nGenes) for LOY and normal nuclei. The boxes represent the 25th percentile, median, and 75th percentile. The whiskers extend to the furthest value that is no more than 1.5 times the inter-quartile range. Significant differences between LOY and non-LOY groups were assessed using the non-parametric Wilcoxon test, resulting in the p-values shown for each comparison. All tests were two-sided. ns P>0.05, *P < 0.05, **P < 0.01, ***P < 0.001.

We and others^13, 14^ have found nuclei with fewer UMI counts and greater expression sparsity have an increased probability of false-positive LOY calls through random dropout of all Y chromosome genes^13^. To confirm that LOY estimates were not a result of low sequencing depth we classified LOY using increasing minimum UMI thresholds (**Supplementary Fig. 15a**). In both the EC and SSC samples, as the minimum UMI threshold was increased from 1000 to 6000, microglia LOY frequency remained stable, while in astrocyte and neuronal populations LOY frequencies approached 0. Furthermore, in astrocyte and neuronal populations, LOY nuclei have significantly lower mean UMI and feature counts than non-LOY/normal nuclei (**Supplementary Fig. 15b-c**), while in microglia these attributes are not significantly different between the groups. Together, this suggests in the ‘Smith’ dataset that we are classifying true LOY nuclei in the microglia, whereas in astrocyte and neuronal populations LOY are more likely to be a result of technical and stochastic factors.

Next, we tested for subtype enrichment of LOY across our seven microglia datasets. We hypothesized that cycling and activated microglia would be most prone to LOY accumulation, however we did not observe substantial differences between subtypes (**Supplementary Table 9**). We were also interested if LOY was associated with infiltrating monocytes. During injury and neurodegeneration, CCR2+ peripheral monocytes can enter the CNS and contribute to neuroinflammation^27^. These cells are commonly identified using markers including CCR2, VCAN and FCN1. In our combined dataset, microglia with monocyte markers were rare and did not show elevated LOY frequencies (**Extended Data Fig. 2**; **Table 2**). However, additional studies specifically designed to investigate LOY in microglia will be required to better elucidate these questions.

**Table 2.** Loss of Y frequency in cells with infiltrating monocyte markers. **Top**, Loss of Y frequency of cells with and without expressed monocyte markers. Microglia and CAMs from each dataset were included. LOY frequency was included if total available cells/nuclei was > 20. **Bottom**, Total cell/nuclei sample size available for each group.

| LOY frequency % |  |  |  |  |  |  |
| --- | --- | --- | --- | --- | --- | --- |
| Cohort | CCR2+ | CCR2- | VCAN+ | VCAN- | FCN1+ | FCN- |
| GSE137444 | NA | 3.20% | NA | 3.20% | 3.70% | 3.20% |
| GSE160936 | 12.80% | 15.80% | 11.80% | 15.90% | 8.60% | 15.80% |
| syn12514624 | NA | 22.80% | NA | 22.70% | NA | 22.70% |
| GSE148822 | NA | 4.20% | 4.20% | 4% | NA | 3.90% |
| GSE178265 | 2% | 3.40% | 2.40% | 3.50% | NA | 3.40% |
| GSE174367 | NA | 5.40% | 7.50% | 5.40% | NA | 5.40% |
| GSE174332 | NA | 3.90% | NA | 3.70% | NA | 3.80% |

| # of cells/nuclei |  |  |  |  |  |  |
| --- | --- | --- | --- | --- | --- | --- |
| Cohort | CCR2+ cells | CCR2- cells | VCAN+ cells | VCAN- cells | FCN1+ cells | FCN- cells |
| GSE137444 | 6 | 7266 | 16 | 7256 | 27 | 7245 |
| GSE160936 | 39 | 12100 | 337 | 11802 | 46 | 12093 |
| syn12514624 | 3 | 2283 | 7 | 2279 | 10 | 2276 |
| GSE148822 | 2 | 1163 | 75 | 1090 | 5 | 1160 |
| GSE178265 | 50 | 7699 | 605 | 7144 | 11 | 7738 |
| GSE174367 | 0 | 2191 | 40 | 2151 | 2 | 2189 |
| GSE174332 | 2 | 856 | 24 | 834 | 3 | 855 |

## Discussion

Mosaic loss of chromosome Y (LOY) has been associated with age-related degenerative CNS diseases^7, 8, 13, 15, 16^, however its incidence and mechanisms in the brain are not well understood. To better characterize LOY-affected CNS cell-types and to inform future LOY studies, we present an extensive analysis of LOY in the brain using single-cell and single-nuclei transcriptomes. Importantly, we provide evidence that amongst CNS cell types, LOY occurs most frequently in microglia (7.79%, *n*=41,949) and causes a transcriptomic signature significantly overlapping those found in peripheral leukocytes with LOY (*P*<0.01). Moreover, 6 of 7 donors with significantly downregulated PAR expression (FDR<0.05) were diagnosed with neurodegenerative disease (AD, PD) or mild cognitive impairment. Together our data indicates LOY occurs in microglia and results in the dysregulation of over 200 genes with roles in aging, glioma biology and inflammation that occur in multiple regions of the brain. Also, we did not find evidence of LOY deriving from infiltrating CCR2+ monocytes, suggesting LOY-causing missegregation events are likely to occur locally in the CNS. Although future studies are required to replicate mLATEs and characterize their roles in disease physiology, we postulate somatic LOY accumulation in the microglia could represent an additional process in age-related dysfunction leading to chronic inflammation and neurodegeneration.

The importance of microglia in age-related dysfunction neurodegenerative processes is rapidly emerging^35, 45^. Age-related alterations in gene expression lead to dystrophic microglia that are less ramified^46^, have a reduced ability to phagocytose debris^47^ and produce greater amounts of proinflammatory cytokines^48, 49^. Further, microglia are long-lived cells^18^ that are derived from EMPs, not hematopoietic stem-cells (HPSC), that turnover locally and are largely isolated from the periphery^21, 23^.

These properties make microglia prone to mutation and selective pressures just as haematopoietic lineages. Moreover, during the progression of many neuropathologies, resident microglia rapidly proliferate, greatly expanding the population and providing an increased opportunity for missegregation errors leading to Y loss^50^. Similar processes occur in aged microglia where a hallmark, “primed”, proinflammatory profile develops resulting in heightened proliferation^51, 52^. Both mechanisms could explain elevated LOY occurrence in microglia. At the same time, spontaneous, somatic mutations in microglia - made increasingly likely by age-related genomic instability – could directly contribute to the pathogenesis of neurodegenerative disease^53^. For example, an induced somatic BRAF(V600E) mutation in murine HPSCs leads to hematological malignancy, but when the same mutation is induced in yolk-sac EMPs it results in a late-onset neurodegenerative disorder^53^. It appears as microglia turnover the mosaic population of BRAF(V600E)-affected, dysfunctional microglia outcompete the normal microglia leading to dystrophy, chronic inflammation, and ultimately neurodegeneration. Further research is required to provide evidence that a similar mechanism leads to LOY selection in microglia, but microglia-specific lineage properties afford plausibility.

Several dysregulated genes in LOY microglia could lead to pathological CNS conditions through processes including immune dysfunction, disruption of homeostatic function, and dysregulated proliferation. Some potential candidates include PAR genes such as Colony Stimulating Factor 2 Receptor Subunit Alpha (CSF2RA) and CD99, which are downregulated in 7 and 4 donor LOY microglia populations, respectively. Within our dataset, microglia, and CNS-associated macrophages (CAM) uniquely express CSF2RA (**Supplementary Fig. 16b**), a receptor subunit for colony-stimulating factor-2 (CSF2). CSF1 and CSF2 signaling is important for brain development and maintenance of microglia homeostasis^54^, and is dysregulated in AD and after brain injury. Imbalance of CSF1R-CSF2 ratios, through CSF2RA deficiency could contribute to chronic inflammation and/or senescence leading to neurodegenerative conditions^54^. CD99 is a cell-surface glycoprotein showing expression in microglia, endothelial cells, pericytes, astrocytes and CAMs (**Supplementary Fig. 16b**). CD99 has multifaceted functions in leukocyte cell adhesion, transendothelial migration, MHC class I transport, and B-cell apoptosis^55^. CD99 also shows complex contradicting roles in various cancers, acting as both a oncosuppressor (osteosarcoma, Hodgkin’s Lymphoma) and oncogene (Ewing sarcoma, malignantglioma)^56^. Adding to the complexity, CD99 presents two specific isoforms with differing roles and cell-type dependent expression^57^. Single-cell evidence shows CD99 surface protein abundance is reduced in leukocyte LOY cells, establishing a link between mRNA and protein dysregulation^14^. Further investigation into the functions of CD99 in microglia are necessary to better understand its potential role in LOY-related processes.

Although historically considered a “functional wasteland” ^58^, recent investigation of the MSY has found several genes that if lost could contribute to cancer and disease development. For example, KDM5D and UTY are histone H3 remodellers, regarded as tumor suppressors in prostate cancer and clear cell renal cell carcinoma^59^. In prostate cancer, loss of KDM5D expression and subsequent H3K4me dysregulation has been associated with expediated cell cycle and mitotic entry^60^. UTY is well expressed in microglia, and similar processes could occur in the CNS (**Supplementary Fig. 16a**). Y-linked lncRNA LINC00278 is also of interest. LINC00278 expression is elevated in microglia and through the AR signaling pathway affects progression of esophageal squamous cell carcinoma^61^.

Our study also highlights 188 autosomal genes and 6 chrX genes dysregulated in LOY microglia that through pleiotropy could plausibly link LOY occurrence with immune and CNS dysfunction. For instance, ROBO1, a major receptor in the SLIT/ROBO signalling pathway, was upregulated in microglia populations from 5 subjects (FDR < 0.05). In addition to important roles in neural development, angiogenesis, metastasis and inflammatory cell chemotaxis, SLIT/ROBO signaling displays contradicting tumor promoting and suppressing properties in various cancers^62–64^. In low-grade glioma (LGG) and glioblastomas (GBM) ROBO1 and SLIT2 are upregulated^65^ and are associated with PI3K-γ activation, poor survival, and resistance to inhibitor therapy^65^. Despite, intriguing roles in glioma metastasis, the links to Y chromosome loss are unknown. However, bulk RNA sequencing in individuals with sex chromosome disorders provides additional evidence that ROBO1 expression is associated with MSY/PAR expression. Compared to controls, ROBO1 is significantly upregulated in Turner syndrome (XO) leukocytes and downregulated in Klinefelter’s syndrome (XXY) (*P*<0.0001, **Supplementary Fig. 17**)^66^. Similar patterns are observed for other upregulated mLATEs including RBFOX2, DOCK1, TPTEP1 and TMTC2 (*P*<0.001) suggesting a PAR dose-dependent transcriptional mechanism. Across all tissues, ROBO1 expression is positively correlated with 117 genes that include RBFOX2, DOCK1, and NAV3 (*P*<0.05; Bonferroni)^67^. Further investigation into PAR associated transcriptional effects are required to uncover mechanisms and if they are associated with disease.

Accompanying the biological findings in this study, technical improvements regarding LOY detection using scRNAseq were made. Mainly, we found that including introns for UMI counting significantly increased detected genes (and MSY genes) with minimal impact on clustering. In 10X Chromium 3’ single-cell assays, to uniquely barcode each individual RNA molecule, poly(dT) primers are designed to prime poly-adenylated (poly-A) mRNA tails^68^. However, poly(dT) primers can also prime internal, often intronic, (poly-A) tracts, resulting in intronic UMIs from mature RNA^68^. Because intronic UMIs are thought to be derived from a biological molecule and are correlated with exonic UMI counts, they represent a valuable information source that improves transcriptional sensitivity when calling LOY. We recommend including introns when attempting to call structural mutations such as LOY using single-cell RNA data.

Although we provide a valuable exploratory analysis of LOY in the brain, there are several limitations with our study. First, because of the biological and technical heterogeneity of our dataset, donor and cell-type sample size was often underpowered when testing significance between LOY and age, and neurodegenerative disorders. Although we found directionally consistent associations between LOY and AD in microglia, the effect of age could not be reliably adjusted for. Single-cell studies specifically designed to test the correlation between LOY and AD in microglia are required. Second, estimating structural mutations such as LOY using the transcriptome is highly variable and prone to several biases that differ between cell-types, individuals, and sequencing batches. The MSY region used to classify LOY only contains ∼15 genes that are commonly detected using scRNAseq, and only 10 that are well expressed (**Supplementary Fig. 16a**). Additionally, sequencing depth per cell varies significantly between experiments, batches, and cell-types, adding to the variability of MSY information. To address this issue, we filtered out cell-type populations in each dataset with insufficient MSY expression (<250% sum MSY pct expressed), and total UMI counts (<3000 UMI). However, transcriptional variability and sparsity remain a confounder that limits the ability to reliably characterize LOY in many datasets. For example, before thresholding, the high-quality ‘Gerrits’ dataset provided 130,936 male single-cell transcriptomes from the occipital cortex, including 7 donors (2 AD), and 39,094 LOY from microglia. However, after filtering only 27,270 nuclei remained. Sequencing depth, MSY expression and PAR expression was shallow, making LOY detection and analysis difficult despite large cell numbers. Similar problems occurred with many other datasets, making LOY detection using LOY difficult. Mattisson *et al*.^14^ have also noted the issues of transcriptional variability with scRNAseq and found that cell-surface proteins are significantly more reliable for characterizing LOY. For example, when measuring CD99 abundance in LOY populations of six leukocyte cell-types, CD99 protein abundance was significantly downregulated in all 6, while RNAseq only found significant CD99 downregulation in 2 cell-types (CD14+ monocytes and NK cells)^14^. Protein abundance is much less sparse than RNA and provides a more reliable LOY readout. To collect the most reliable information on LOY-specific transcriptional effects, future LOY studies should utilize multiomic single-cell technologies such as CITE-seq, or ATAC+RNAseq as we showed above. CITE-seq has been applied to CNS immune cells in mice^69^, and similar studies in aging men would be valuable for further characterizing LOY in the CNS.

In conclusion, using single-cell transcriptomes we present the first evidence of LOY affecting microglia in the brain, disproportionately affecting elderly donors diagnosed with neurodegenerative disorders. Our results show microglial LOY affects the expression of hundreds of autosomal genes with diverse functions that require additional experimental investigation. Finally, we believe LOY in the microglia could represent an additional, understudied biological process that could alter microglia phenotypes and play a role in neurodegeneration.

## METHODS

### Single-cell/nuclei dataset curation and selection

Single-cell transcriptome data used in this study was acquired from both public and controlled sources (**Supplementary Table 1**). Publicly available single-cell/nuclei RNA-seq datasets were downloaded from the Gene Expression Omnibus (GEO), Human Cell Atlas (HCA) and 10x Genomics dataset page (https://www.10xgenomics.com/resources/datasets). Controlled access data was downloaded from Synapse through the AMP-AD Knowledge Portal. We searched for datasets that met the following criteria: (1) available raw FASTQ files (single-cell) or unnormalized count matrix (single-nuclei); (2) were generated using the 10x Genomics Chromium platform; and (3) profiled human samples from primary human brain tissue. By using data produced via 10x Chromium library preparation we reduced undesired technical variation. In total, we collected 21 datasets across a diverse set of brain regions that totalled 763,410 male cells/nuclei from 253 male individuals (median age: 71.5; range age: 0-96). Both male and female cells were included for initial clustering and annotation.

### Single-cell and single-nuclei RNAseq processing

#### Generating expression matrices (single-cell)

Beginning with FASTQ files, all single-cell RNAseq datasets used in this study (*n*=4) were processed through the same bioinformatic pipeline (**Supplementary Fig. 1**). For each dataset, alignment of reads and cell barcode filtering were performed using Cellranger 5.0.0 (10x Genomics). Reads were aligned to the GRCh38 2020-A reference (10x Genomics) using Cellranger *count* in intron retaining mode. To include intronic reads, we used the ‘ –include-introns’ flag. This flag considers quality, confidently mapped intronic reads as candidates for UMI counting. Although the utilization of intronic reads is commonly used for samples enriched with pre-mRNA (i.e. snRNAseq experiments) we observed improved gene detection sensitivity with minimal impact on clustering. Similar findings were made here^68^. Our data shows when performing aneuploidy estimation using scRNAseq, increased transcriptional sensitivity and greater cell/nuclei counts improves the power of LOY DE tests. Sample-specific UMI count matrices output from *count* were aggregated using cellranger *aggr* without read count normalization.

#### Generating expression matrices (single-nuclei)

Most published single-nuclei datasets are generated using intronic information and therefore processing through our local pipeline did not provide noticeable transcriptional sensitivity benefits. Furthermore, because of patient privacy concerns several single-nuclei datasets had restricted access to raw FASTQ files. To save time and computational resources we used publicly available expression matrices for most single-nuclei datasets. Processing details for each dataset is available in **Supplementary Table 1.**

#### Quality control, filtering, and clustering

Gene expression matrices were imported into R (v4.0.2) for further analysis using Seurat (v4.0.2)^70^. Filters were applied to retain cells/nuclei with adequate gene representation (>1000 genes), unique molecular identifier counts (>1500 UMI), mitochondrial UMI percentage (<15%), and ribosomal UMI percentage (<45%). These filters were applied to discard potential empty droplets and apoptotic cells. scDblFinder (v1.6.0)^71^ was applied using default settings to identify and remove putative doublets. Normalization, dimension reduction, clustering and annotation were performed using Seurat (**Supplementary Methods**). If systemic biases were observed, batch correction was applied using Harmony (v1.0.0)^72^.

In-depth microglia subcluster analysis was completed for 7 datasets including syn12514624, GSE148822, GSE137444 and GSE160936, GSE178265, GSE174367 and GSE174332 (**Supplementary Fig. 6-12; Supplementary Table 9**). Each included dataset was subset for microglia. Individual samples were integrated using canonical correlation analysis (CCA). *Seurat* CCA integration was used instead of *Harmony* when analyzing in-depth subsets and clustering patterns. All other filtering and processing were completed as forementioned (**Supplementary Methods**). To annotate biologically relevant subclusters within microglia we complied gene sets from several published studies and calculated module scores via the Seurat “AddModuleScore” function (default settings). All gene sets used in the study are provided in **Supplementary Table 10**.

#### Loss of Y classification

Our method of determining LOY classification from each single-cell transcriptome was adapted from previous studies^1, 13^. Most male cells commonly express several genes located in the male-specific region (MSY) of the Y chromosome (GRCh38:Y:2,781,480-56,887,902). In each dataset, cells were classified as LOY if they lacked expression of all commonly expressed genes residing in the MSY, which were defined as MSY genes with >0.05 normalized expression and/or expressed in >5% of cells. Furthermore, in datasets with subjects from both sexes, MSY genes with comparable normalized expression between male and female cells/nuclei were removed (> 0.2 ratio). This was done to exclude chrY features whose expression may be contaminated by homologous genes located on chrX. Ultimately, cells were labelled LOY if they lacked detection of all expressed MSY genes and were labelled normal/non-LOY if they did not. Tools used to classify LOY can be accessed at https://github.com/michaelcvermeulen/scLOYtools.

#### Loss of Y proportion

We identified LOY frequencies by comparing the number of LOY cells to total cells within each cell-type, within each subject. Additional filters were applied to limit variability and technical biases. For LOY frequency analysis we used stricter QC threshold (nUMI > 3000 and nGene > 1000) to limit false-positive LOY calls. Cell-type populations within each subject with <75 cells/nuclei were removed to limit variability. Also, preliminary analysis found that in several sequencing experiments, some cell-type populations lacked sufficient MSY detection, often resulting in LOY frequency overestimation. To limit this effect, we filtered populations based on summed MSY percent expressed. This metric was calculated in non-LOY cells and represents the summed, percent expressed value of all MSY genes, giving a measure of how common MSY expression was in non-LOY cells. Cell-type populations with less than 250% summed MSY expression were removed (**Supplementary Methods**).

#### Loss of Y differential gene expression

We identified LOY-associated transcriptional effects (LATE) by performing DE tests between LOY and non-LOY populations. To limit false positive LOY classification influenced by high gene dropout and low information, minimum UMI (>1500) and gene (>1000) filters were applied. DE analysis was performed for each cell-type using MAST algorithm via the Seurat *FindMarkers* function using full datasets (tissue-specific) and individual subjects. When performing DE on a full dataset, latent variables including sample, percent mitochondrial RNA, nUMI and nGenes were added to the hurdle-model and regressed out. For tests on single subjects, we used tissue of origin, percent mitochondrial RNA, nUMI and nGenes as latent variables. Genes were included if they were expressed in >5% of cells/nuclei.

#### Gene set enrichment analysis (GSEA)

All genomic region, pathway, and GO term overrepresentation for differential expressed genes was performed using Molecular Signatures Database (MSigDB) gene sets^73^ and the hypergeometric test provided by the hypeR package^74^. DE genes with FDR<0.05 were used and significance for all GSEA tests was declared using FDR<0.05. To determine PAR region enrichment the c1.all.v7.4.symbols.gmt file was edited to include PAR1 and PAR2 as independent cytogenic bands. In all datasets that were analyzed, PAR2 lacked regular expression and therefore PAR1 was used and considered PAR. The c2.cp.v7.4.symbols.gmt set was used for pathway enrichment and c5.go.v7.4.symbols.gmt was used for GO term enrichment.

#### Single nuclei multiome ATAC and gene expression analysis

The multimodal LOY analysis was performed on sample data provided by 10x Genomics (https://www.10xgenomics.com/resources/datasets/fresh-frozen-lymph-node-with-b-cell-lymphoma-14-k-sorted-nuclei-1-standard-2-0-0). The dataset contains joint ATACseq and RNAseq for 14000 nuclei from an intra-abdominal lymph node tumor from a male patient diagnosed with non-Hodgkin lymphoma. Filtered feature barcode matrices (HDF5) and fragment files were downloaded from 10X Genomics and processed using both Seurat (v4.0.2) ^70^ and Signac (v1.2.1) ^75^, following the Signac 10X scATAC vignette (https://satijalab.org/signac/articles/pbmc_vignette.html). Briefly, ATAC data was normalized using TF-IDF and features in the 75^th^ quartile were considered top features. Peaks were identified using Cellranger and peaks within blacklist regions were removed (blacklist_hg38_unified). Gene expression pre-processing was performed as above, and dimension reduction and clustering were performed solely using the transcriptome. HG38 gene annotations from standard chromosomes were added from EnsDb.Hsapiens.v86. To quantify the relative activity of each gene based on chromatin accessibility we used the *GeneActivity* function (Signac). Within each cell *GeneActivity* extracts gene coordinates (extended 2kb upstream to include promoter) and counts fragments in each region. The gene activity matrix was log normalized and scaled then added as an assay to the Seurat object. For each cell, MSY gene activities were calculated by taking the mean normalized gene activity of all detected MSY genes (RPS4Y1, ZFY, TBL1Y, USP9Y, DDX3Y, UTY, TMSB4Y, NLGN4Y, KDM5D, EIF1AY).

### Other statistical analyses

All statistical analyses were performed using R (v4.0.4). Fisher’s exact test used determine overlap significance between gene sets, was performed using the GeneOverlap package ^76^.

Non-parametric Wilcoxon tests were performed using the wilcoxon.test function from the R stats package. Global gene co-expression analysis was performed using correlationAnalyzeR (Bonferroni <0.05) ^67^. Plotting was done using a combination of Seurat, Signac, ggpubr, ggplot2 and dittoSeq packages.

### Author Contributions

MV and SM conceived the project. MV carried out analysis and experiments, and wrote the first draft of the paper. SM, TYP, RP provided feedback. All authors contributed to writing and finalizing the draft.

---

**Extended Data Fig 1.**
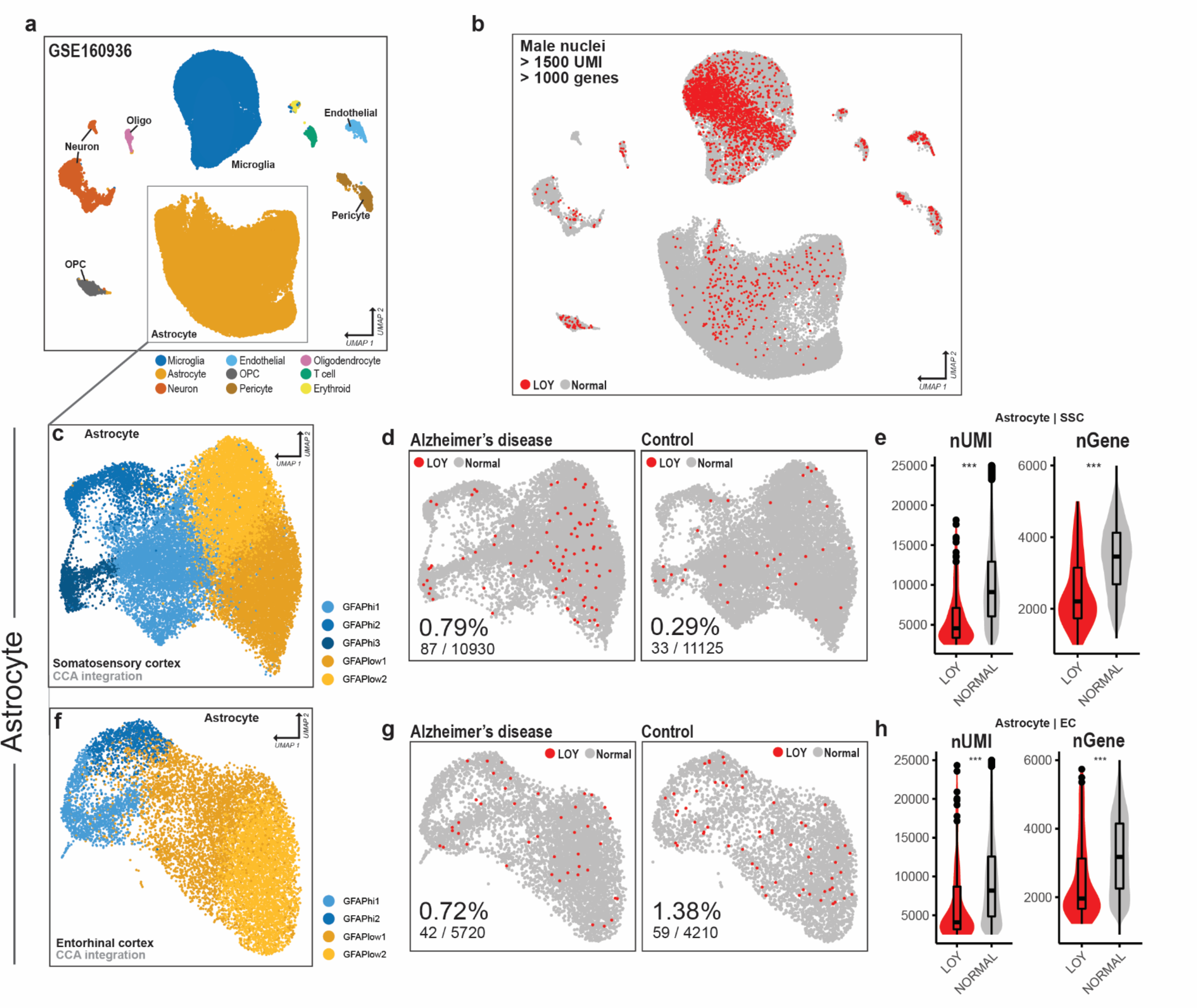
LOY frequencies in astrocytes across the EC and SSC (GSE160936). **a**, UMAP reduction and cell-type annotations of 48,748 single-nuclei brain transcriptomes from GSE160936. Markers used to annotate clusters include C3 and CSF1R (microglia), AQP4 and GFAP (astrocyte), RBFOX3 and GAD1 (neuron), and MBP (oligodendrocyte). All marker genes used to annotate clusters are available in Supplementary Fig. 9. **b**, UMAP plot showing LOY classification of each male nucleus (nUMI > 1500 and nGene > 1000). LOY nuclei (red) and non-LOY/normal nuclei (grey). Astrocyte nuclei from SSC (**c**) and EC (**f**) were isolated, CCA integrated, and clustered. **c**,**f**, Subclusters were annotated using high (yellows) and low (blues) GFAP expression. **d,g**, UMAP reductions of SSC and EC split by AD diagnosis; Nuclei are colored by LOY (red) and non-LOY/normal (grey). LOY nuclei frequencies are provided at the bottom of each UMAP plot. **e,h**, Violin boxplots showing UMI counts (nUMI) and detected genes (nGenes) for LOY and normal nuclei. The boxes represent the 25th percentile, median, and 75th percentile. The whiskers extend to the furthest value that is no more than 1.5 times the inter-quartile range. Significant differences between LOY and non-LOY groups were assessed using the non-parametric Wilcoxon test, resulting in the *p*-values shown for each comparison. All tests were two-sided. *ns P*>0.05, \**P* < 0.05, \*\**P* < 0.01, \*\*\**P* < 0.001.

**Extended Data Fig. 2.**
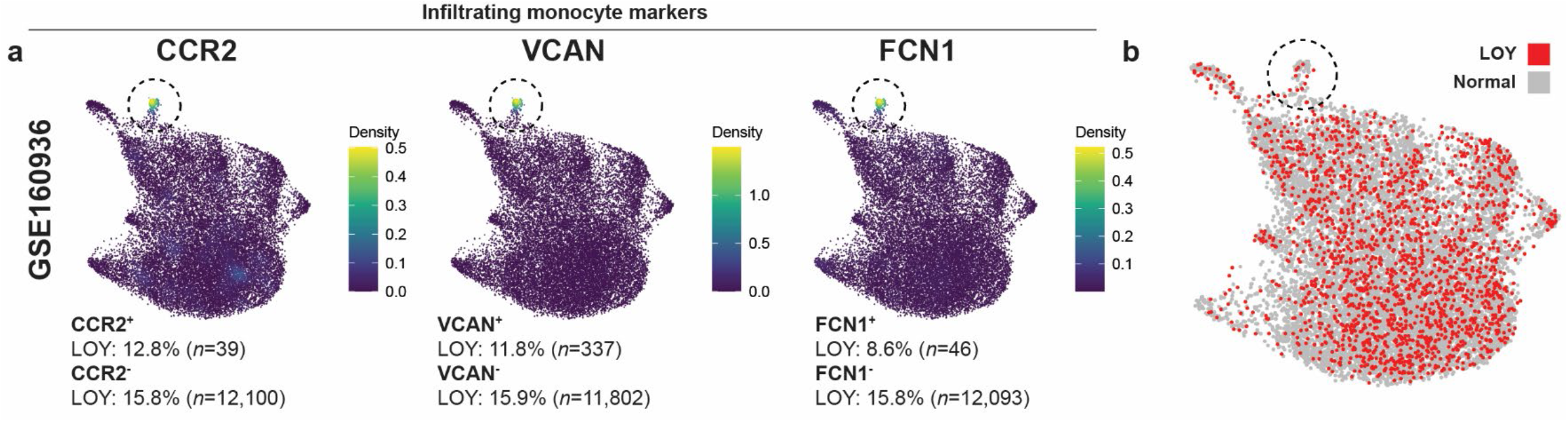
Infiltrating monocytes do not harbor disproportionate LOY frequencies. **a**, Three microglia UMAP plots displaying kernel gene-weighted density estimation for three known markers of infiltrating monocytes in the CNS (CCR2, VCAN and FCN1) from dataset GSE160936. Monocyte expression was confined to one small cluster and was detected in 674 nuclei (2.7%; male and female). Text below each plot displays LOY proportions for male nuclei with and without detected CCR2, VCAN and FCN1. Total sample size is provided in parentheses. Nuclei were included if they contained > 3000 nUMI and >1000 nGenes. **b**, UMAP reduction displaying 12,139 male microglia, monocyte, and CNS associated macrophage (CAM) nuclei from GSE160936 (from the somatosensory cortex and entorhinal cortex). Each point represents an individual nucleus and is colored by LOY (red) and non-LOY (grey).

